# Pathway centrality in protein interaction networks identifies functional mediators of pulmonary disease

**DOI:** 10.1101/171942

**Authors:** Jisoo Park, Benjamin J. Hescott, Donna K. Slonim

**Author notes:** Correspondence (D.K.S.) (J.P.).

## Abstract

Identification of functional pathways mediating molecular responses may lead to better understanding of disease processes and suggest new therapeutic approaches. We introduce a method to detect such mediating functions using topological properties of protein-protein interaction networks. We introduce the concept of pathway centrality, a measure of communication between disease genes and differentially expressed genes. We find mediating pathways for three pulmonary diseases (asthma; bronchopulmonary dysplasia (BPD); and chronic obstructive pulmonary disease (COPD)) using pathway centrality. Mediating pathways shared by all three pulmonary disorders heavily favor inflammatory or immune responses and include specific pathways such as cytokine production, NF Kappa B, and JAK/STAT signaling. Disease-specific mediators, such as insulin signaling in BPD or homeostasis in COPD, are also highlighted. We support our findings, some of which suggest new treatment approaches, both with anecdotal evidence from the literature and via systematic evaluation using genetic interactions.

## Introduction

It has long been noted that genes with variants implicated in disease are not necessarily differentially expressed, and that differential expression does not easily lead to the discovery of disease genes (Hudson et al., 2012). In many cases, differential expression predominantly reflects the tissue-specific consequences or downstream effects of a disease-causing process that integrates complex genetic and environmental responses. This makes differentially expressed genes useful as diagnostic markers, but often poor as therapeutic targets (Fox et al., 2011). Conversely, functional analysis of causal disease genes, whether identified through highly specific cell or animal studies or systematically via GWAS, often won’t fully explain how these downstream responses occur (Delude, 2015). However, we know that other functional pathways may mediate the expression responses we see. We hypothesize that finding these can provide new insights into disease processes and suggest novel therapeutic approaches. Here, we test this hypothesis by examining three different pulmonary disorders from different life stages to identify common and unique mediating mechanisms in airway disease.

We start by considering the roles that proteins corresponding to disease genes or differentially expressed genes play in protein-protein interaction networks. We postulate that the functional pathways mediating disease response will disproportionately reflect communication between these two sets of proteins, and conversely, that if we look for such “central” pathways we will find mediators. To identify putative mediating pathways in disease response, we first introduce a generalized notion of network centrality called *pathway centrality*, which combines a variation of *betweenness* with the concept of *group centrality* (Everett and Borgatti, 1999). Node betweenness characterizes how many of the shortest paths between all pairs of nodes pass through a given node (Abman et al., 1987, Yu et al., 2007). Our variation of betweenness counts only the shortest paths between disease genes and differentially expressed genes passing through a given node. The notion of group centrality averages the betweenness scores of a set of nodes to measure the centrality of the whole set. Therefore, our pathway centrality score is the average modified betweenness score of a set of genes participating in the given pathway.

There have been many other approaches to defining centrality in protein interaction networks (Mason and Verwoerd, 2007). Some have focused on modeling communication between nodes by network or graph flows (Freeman et al., 1991). Using network flows to calculate network distance has proven helpful for protein function prediction (Nabieva et al., 2005), and has been compared favorably to betweenness centrality for integrating function prediction with contextual data (Missiuro et al., 2009). However, this approach has not yet been applied to group centrality in protein interaction networks. Viewing group centrality as an optimization problem has also been used to discover new groups of important nodes in networks (Erdos, 2015), but not for identifying functional gene sets playing a pivotal role.

Most relevant to our efforts is a collection of prior results linking expression quantitative trait loci (eQTLs) to differentially expressed genes via protein-protein, protein-DNA, and phosphorylation networks. These studies were initially intended to find the exact causal gene in a linked locus using pathway information. One early approach (Tu et al., 2006) uses a node’s accessibility in random walks to prioritize causal genes and identify regulatory paths through the network. Slightly later work (Suthram et al., 2008, Kim et al., 2011) models the integrated network as an electrical circuit, identifying putative causal genes by the flow of current through the network. Kim, *et al.* explicitly extended the eQTL-target work to a disease context (Kim et al., 2011). Specifically, by linking copy number variations, through eQTL mapping, to differentially expressed genes, they identified likely causal mutations in glioblastoma multiforme. Similarly, Yeger-Lotem, *et al.* used integrated protein-protein and protein-DNA interaction networks to identify proteins and genes on high-probability paths between “genetic hits” and transcriptionally regulated genes (Yeger-Lotem et al., 2009). In this work, which focuses on a number of cellular perturbations in yeast, the emphasis is on finding individual proteins on these paths and exploring the functions they perform.

These efforts are related to ours in the sense that they examine information flow between genes linked to disease and differentially expressed genes. Specifically, our disease genes come from a compilation of sources that includes, but is not limited to, GWAS data. However, here our focus is limited to the disease-related pathways, and our aim is to identify underlying biological functions that mediate cellular response in disease, rather than to identify causal mutations. Given a disease *d*, a set D of known disease genes for *d*, and a set E of genes differentially expressed in relevant tissues as a consequence of disease *d*, we search for gene sets M that significantly participate in passing signals from D to E (Figure 1). We calculate the pathway centrality (PC) score for a candidate pathway S considering only the paths from nodes in D to nodes in E. We then use permutation tests to assess whether S has higher pathway-centrality than expected (see Methods). For each gene set S, the percentage of such random sets with higher pathway centrality scores than S is reported as *p_cent_*(S), a rough measure of significance.

**Figure 1.**
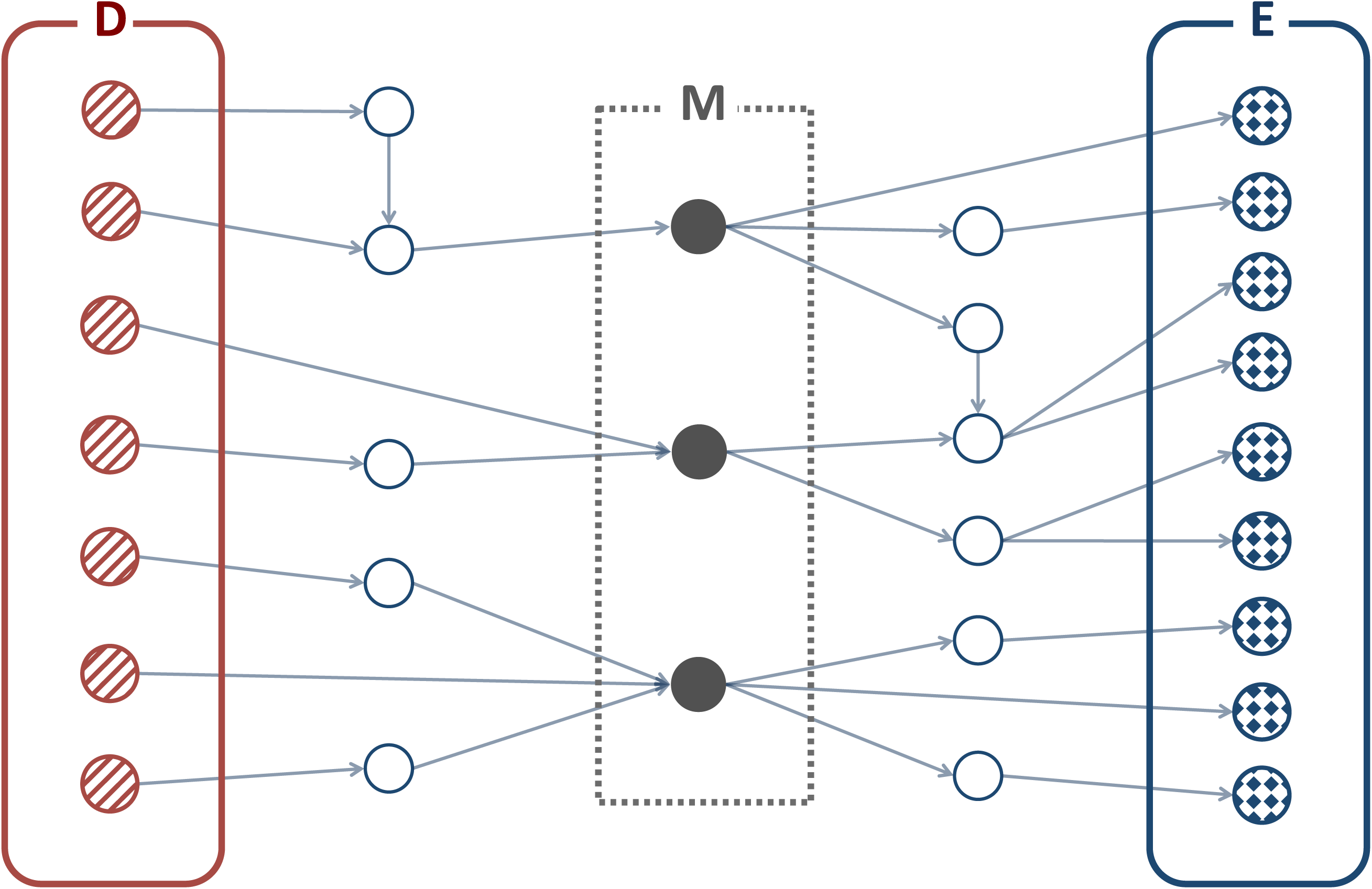
Hypothesized topological property of mediating pathways. Given that D is a set of disease genes and E is a set of differentially expressed genes, mediating pathway genes, M, will participate more significantly in passing signals from D to E.

In this initial study, we apply our novel pathway-centrality method to three pulmonary diseases that primarily affect patients at different stages of life: bronchopulmonary dysplasia (BPD), a neonatal complication of preterm birth; asthma, which is relevant across the lifespan but is becoming increasingly common in children; and chronic obstructive pulmonary disease (COPD), a term that encompasses a number of progressive lung disorders that predominantly affect the elderly (Suzuki et al., 2016). We examine mediating pathways in each disease and look for common pathways relating all three.

One caveat is that pathways with low pathway centrality significance (*p_cent_*) scores for multiple diseases might be highly central in the network structure overall, and could appear to be “significant” for any disease considered. Therefore, as a further control, we applied the method to Down syndrome (DS) data, with the understanding that most functions relevant in pulmonary disorders are probably not that relevant to the etiology of DS, although we agree that there may be some exceptions. Our “disease genes” in this case are simply any genes located on chromosome 21, rather than genes that have been directly implicated in the etiology of DS symptoms.

Supplementary Figure S1 shows the overlaps between the four disease gene sets and the four differentially-expressed gene sets. Although there is some overlap between the sets of disease genes, particularly involving genes involved in inflammation and immune response, overall the disease gene sets are reasonably disjoint and the differentially-expressed gene sets even more so. Thus, common pathways across all three networks are unlikely to have arisen from shared shortest paths between identical sets of genes.

Pathway gene sets used in our experiments come from the Biological Process (BP) terms in the Gene Ontology (Ashburner et al., 2000), pathways in the Kyoto Encyclopedia of Genes and Genomes (KEGG) (Kanehisa et al., 2016), and mammalian phenotypes (MP) from the International Mouse Phenotyping Consortium in the Mouse Genome Database (Bult et al., 2016).

Our results show that pathways involved in inflammation or adaptive immunity, and several related signaling pathways including NFKB, JAK/STAT, and toll-like receptor signaling, are common to all three pulmonary disorders. Disease-specific findings include protein homeostasis pathways in COPD, leukocyte chemotaxis in asthma, and regulation of map kinase activity in BPD. While many of our findings have already been proposed as disease mediators, diagnostics, or targets in previous publications, we discovered several novel mediators that may suggest new therapeutic approaches to these pulmonary disorders.

## Results and Discussion

### Pathway centrality finds mediating pathways and potential drug targets

Pathways with significant pathway centrality in specific pulmonary data sets but not in the Down syndrome data include known disease mediators and potential targets. Full lists of results are available as supplemental data; Table 1 shows the top few pathways implicated in exactly one of the pulmonary disorders. We highlight some of these top pathways here.

**Table 1.** Most-significant GO Biological Processes and KEGG pathways in individual pulmonary disorders. These pathways are the top few with p_cent_< 0.05 in exactly one of the pulmonary disorders and p_cent_ > 0.05 in the DS data. Cells containing p-values below 0.05 are highlighted in boldface. Pathways shown here were selected by criteria discussed in the text; full results are available as supplemental data.

| Pathway | asthma p-value | BPD p-value | COPD p-value | DS p-value |
| --- | --- | --- | --- | --- |
| *GO Biological Process Terms:* |  |  |  |  |
| Regulation of DNA Metabolic Process | **0.0014** | 0.1015 | 0.2034 | 0.3279 |
| Leukocyte Chemotaxis | **0.0021** | 0.3232 | 0.1634 | 0.2219 |
| G1 Phase | **0.0027** | 0.0771 | 0.1375 | 0.087 |
| Leukocyte Migration | **0.0032** | 0.4966 | 0.2705 | 0.3176 |
| Regulation of MAP Kinase Activity | 0.1779 | **0.0035** | 0.0502 | 0.0874 |
| Ras Protein Signal Transduction | 0.0995 | **0.0049** | 0.0854 | 0.1305 |
| Homeostatic Process | 0.156 | 0.1313 | **0.0004** | 0.362 |
| Nuclear Transport | 0.2514 | 0.056 | **0.0018** | 0.089 |
| Nucleocytoplasmic Transport | 0.2455 | 0.0524 | **0.002** | 0.0872 |
| Chemical Homeostasis | 0.2365 | 0.1249 | **0.002** | 0.6049 |
| *KEGG Pathways:* |  |  |  |  |
| FC Gamma R Mediated Phagocytosis | **0.0044** | 0.18 | 0.111 | 0.3449 |
| Tight Junction | **0.0128** | 0.0564 | 0.135 | 0.0927 |
| Regulation of Actin Cytoskeleton | 0.2283 | **0.0026** | 0.103 | 0.2323 |
| Antigen Processing and Presentation | 0.2898 | **0.0053** | 0.0932 | 0.1658 |
| Insulin Signaling Pathway | 0.3315 | **0.0195** | 0.0966 | 0.2575 |
| Cell Adhesion Molecules (CAMs) | 0.7917 | 0.1658 | **0.0198** | 0.3768 |
| RIG I Like Receptor Signaling Pathway | 0.0726 | 0.1353 | **0.0235** | 0.1229 |
| Viral Myocarditis | 0.3229 | 0.0825 | **0.0391** | 0.3046 |

Highly significant in asthma is the BP gene set *leukocyte chemotaxis.* Neutrophil chemotaxis velocity has been suggested as a biomarker for asthma and incorporated into a diagnostic test to distinguish asthma from nonasthmatic allergic rhinitis (Sackmann et al., 2014). The KEGG gene set *cell adhesion molecules* tops the list in COPD, pointing at similar processes, although the only genes shared between these two gene sets are *ITGB2* and *ITGA9.* Prior work suggests that adhesion molecules also play a significant role in the pathogenesis of COPD (Ishii, 1999), and that regulation of adhesion may be a new therapeutic approach for both COPD and asthma (Chen et al., 2008, Woodside and Vanderslice, 2008). The KEGG pathway *tight junction* is also implicated in asthma. Although adhesion and leukocyte chemotaxis are important to all three disorders (Faura Tellez et al., 2016, Ramsay et al., 1998, Barnes et al., 2009), the pathway centrality approach highlights different sets of genes mediating these responses. Different pathways have been implicated in recruitment of neutrophils in different contexts in COPD (Barnes et al., 2009), suggesting the plausibility of context-specific variation.

Topping the GO COPD list are two terms relating to homeostasis. Protein homeostatic imbalance has been implicated in the pathogenesis of COPD, although this approach to characterizing the disease has not yet been thoroughly explored (Bodas et al., 2012). Such imbalances have been successfully targeted through small molecules or gene therapies in other diseases such as cystic fibrosis. Addressing the imbalance of critical NFKB-regulated inflammatory proteins has thus been suggested as a possible therapeutic approach in COPD and other airways diseases (Bouchecareilh and Balch, 2012).

Mitogen-activated protein kinases (MAPKs) regulate many developmental and cellular processes (Kim and Choi, 2010), including inflammation and apoptosis. While these pathways are likely involved to some degree in all three pulmonary disorders, the top GO BP gene sets implicated in BPD are *regulation of map kinase activity* and *ras protein signal transduction.* The extracellular signal-related kinases ERK1/2 are major components of the MAPK pathway, which can be activated by the Ras GTP-ase (Kim and Choi, 2010). There is also evidence that ERK/MAPK activation can protect against the negative effects of hyperoxia on alveolar development and lung epithelial cells (Xu et al., 2006, Buckley et al., 2005). Drugs that promote this process have been suggested as potential therapies for BPD (Sakurai et al., 2013).

The KEGG insulin signaling pathway is also implicated as a mediator in BPD. The role of insulin signaling in BPD is unclear, but low neonatal serum IGF-1 levels have been shown to be associated with the later development of BPD (Lofqvist et al., 2012). This evidence is consistent with cell and animal studies showing that IGF-1 signaling, mediating by NFKB, enhances collagen synthesis in fetal lung fibroblasts (Chetty et al., 2006), and that IGF-1 and IGFR1 regulate alveologenesis in neonatal rats (Belcastro et al., 2015). Whether this finding suggests novel strategies for BPD treatment, however, is an open question.

### Commonalities across all three pulmonary disorders implicate immune processes and signaling pathways

When we look for pathways that play a significant role across *all three* pulmonary disorders (Table 2), we are not surprised to find a preponderance of inflammatory and immune processes. GO terms topping the list implicate the adaptive immune response, cytokine production, and pro-inflammatory *NF Kappa B* signaling. Recent research implicates *IKK*-driven *NFKB* activation of inflammation in both COPD and asthma, but suggests that the different components of the system involved in the two diseases could explain their differing pharmacological responses (Gagliardo et al., 2011), and could suggest new avenues for therapy. *NFKB*-mediated inflammation has also been implicated in the pathogenesis of BPD. The increased prevalence of early-onset emphysema in BPD survivors (Wong et al., 2008) supports the hypothesis that an *NFκB*-related inflammatory phenotype predisposes individuals to an increased reaction to environmental airway stress (Park et al., 2014). Links between BPD and asthma through *NFKB1A* promoter polymorphisms have also been identified (Ali et al., 2013).

**Table 2.** All GO Biological Process terms and KEGG pathways, and manually selected Mammalian Phenotype terms, significant (p < 0.05) in all three pulmonary disorders and with p > 0.05 in the DS data. There were 58 such MP terms in total.

| Pathway | asthma p-value | BPD p-value | COPD p-value | DS p-value |
| --- | --- | --- | --- | --- |
| *GO Biological Process Terms:* |  |  |  |  |
| Cell Surface Receptor Linked Signal Transduction | 0.0447 | 0 | 0.0091 | 0.2122 |
| Protein Kinase Cascade | 0.0058 | 0.0011 | 0.0075 | 0.2083 |
| Positive Regulation of Response To Stimulus | 0.0269 | 0.0017 | 0.0043 | 0.1066 |
| Positive Regulation of Immune System Process | 0.0343 | 0.0019 | 0.0059 | 0.1561 |
| Regulation of Immune Response | 0.02 | 0.0025 | 0.0023 | 0.0841 |
| Positive Regulation of Immune Response | 0.0174 | 0.0028 | 0.0024 | 0.0711 |
| Regulation of Immune System Process | 0.0443 | 0.0041 | 0.0173 | 0.2604 |
| Regulation of Response To Stimulus | 0.0383 | 0.0043 | 0.0198 | 0.199 |
| I KappaB Kinase NF KappaB Cascade | 0.0021 | 0.0046 | 0.0115 | 0.1327 |
| Positive Regulation of Multicellular Organismal Process | 0.0487 | 0.005 | 0.0153 | 0.2606 |
| Activation of Immune Response | 0.017 | 0.0074 | 0.0096 | 0.1492 |
| Regulation of Cytokine Production | 0.0144 | 0.019 | 0.0075 | 0.0546 |
| Adaptive Immune Response GO 0002460 | 0.0221 | 0.0283 | 0.0384 | 0.0623 |
| Adaptive Immune Response | 0.025 | 0.0289 | 0.0476 | 0.0697 |
| Protein Polyubiquitination | 0.0133 | 0.0435 | 0.0167 | 0.0683 |
| *KEGG Pathways:* |  |  |  |  |
| Leishmania Infection | 0.0002 | 0.0032 | 0.0098 | 0.1635 |
| JAK STAT Signaling Pathway | 0.0003 | 0.0001 | 0.0097 | 0.1072 |
| B Cell Receptor Signaling Pathway | 0.0003 | 0.0004 | 0.0091 | 0.0662 |
| FC Epsilon RI Signaling Pathway | 0.0004 | 0.0006 | 0.0236 | 0.1027 |
| Natural Killer Cell Mediated Cytotoxicity | 0.0058 | 0.001 | 0.0473 | 0.5846 |
| Chemokine Signaling Pathway | 0.0065 | 0 | 0.0025 | 0.4633 |
| Toll Like Receptor Signaling Pathway | 0.0136 | 0.0094 | 0.0294 | 0.247 |
| Acute Myeloid Leukemia | 0.0175 | 0 | 0.0158 | 0.0637 |
| T Cell Receptor Signaling Pathway | 0.0275 | 0 | 0.0206 | 0.0574 |

| *Selected Mammalian Phenotype Terms:* |  |  |  |  |
| --- | --- | --- | --- | --- |
| Immune System Phenotype | 0 | 0 | 0.0001 | 0.0654 |
| Decreased Bone Resorption | 0.0003 | 0.011 | 0.0175 | 0.1103 |
| Abnormal Circulating Complement Protein Level | 0.0047 | 0.0245 | 0.0063 | 0.2421 |
| Increased Circulating Angiotensinogen Level | 0.0121 | 0.0117 | 0.0172 | 0.1478 |
| Abnormal Circulating Chemokine Level | 0.0176 | 0.015 | 0.018 | 0.1993 |
| Abnormal Airway Responsiveness | 0.0222 | 0.0067 | 0.0372 | 0.556 |
| Abnormal Chemokine Level | 0.0341 | 0.0131 | 0.017 | 0.2805 |
| Abnormal Interleukin-4 Secretion | 0.0427 | 0.0055 | 0.0424 | 0.1368 |

The KEGG and MP gene set collections implicate several more specific pathways, including different types of lymphocyte responses, *JAK*/*STAT* signaling, toll-like receptor signaling, and *FC Epsilon RI* signaling. The *JAK*/*STAT* pathway has been suggested as an asthma target through inhibitors of activating cytokines and receptors (Vale, 2016, Weltman and Karim, 1998). *JAK* pathway inhibitors are in development for a number of inflammatory disorders (O’Shea et al., 2015), and animal models have suggested that targeting this pathway can reduce airway hyperresponsiveness, reflecting the potential for such compounds in both COPD and asthma (Barnes, 2016). The role for *JAK*/*STAT* signaling in bronchopulmonary dysplasia is less clear, but it has been suggested that it plays a role in airway smooth muscle mitogenesis, implicated in both asthma and BPD (Simon et al., 2002), and postulated that it may be an alternative mediator of the oxidative stress response in both diseases (Zhou and Hershenson, 2003). Thus, our work suggests that the effects of *JAK* pathway inhibitors may be worth exploring in pre-clinical models of bronchopulmonary dysplasia.

Toll-like receptor (TLR) signaling, which activates the innate immune response, is another familiar part of the story of airway hyperreactivity and fetal lung development (Petrikin et al., 2010). TLR polymorphisms have been linked to an increased risk of developing BPD (Carrera et al., 2015, Malash et al., 2016), and TLR agonists are already being tested for therapeutic efficacy in asthma (Biggadike et al., 2016). However, the role of this system in COPD is not as clear. Aspects of the innate immune response are often demonstrably suppressed in COPD patients (Shaykhiev and Crystal, 2013), and TLR polymorphisms play a role in disease susceptibility and severity (Apostolou et al., 2016, Yu et al., 2016). Our work therefore also suggests a role for TLR pathways in the diagnosis, stratification, and treatment of COPD.

Finally, the *FC Epsilon RI* signaling pathway raises interesting questions about common elements of these three diseases. Figure 2 shows a subset of this pathway and the network for BPD. The gene *FCER1A* is one of the primary receptors for immunoglobulin E (IgE) (Sharma et al., 2014), the key player in initiating allergic response. The role of allergy in these three disorders, however, is thought to be quite different. The significant impact of IgE response in allergic asthma is well-studied (Gould and Sutton, 2008). Its role in COPD is less clear. Many patients with COPD but no asthma diagnosis have one or more asthma-like symptoms, including atopy, and there are suggestions that allergic response plays a role in severity for a subset of those with COPD (Suzuki et al., 2016).

**Figure 2.**
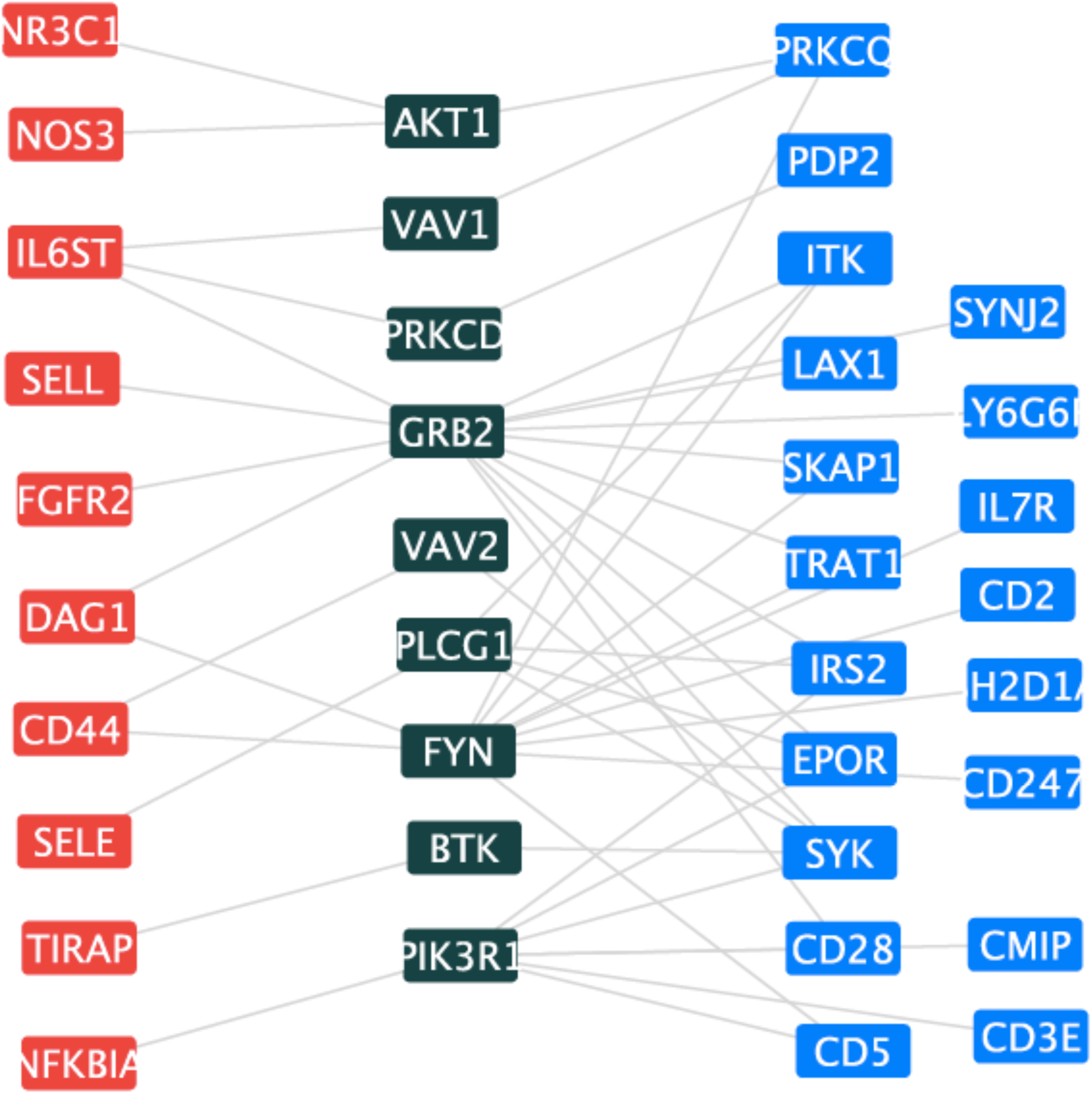
Topology of BPD-related genes and the *FC Epsilon RI* signaling pathway. One of the significant mediating pathways for Bronchopulmonary Dysplasia (Carrera et al., 2015) is the KEGG *FC Epsilon Rl* signaling pathway. Here, pathway genes are colored dark green, mediating pathway genes are in between BPD disease genes (red) and differentially expressed genes (blue). The VAV proteins and BTK, for example, are known to be involved in lymphocyte development and activation.

In contrast, most evidence suggests that bronchopulmonary dysplasia is *not* linked to the development of allergies or to IgE response (Kwinta et al., 2013). Indeed preterm birth has been shown to correlate with a *decreased* risk of atopy (Siltanen et al., 2001), although with an increased risk of asthma and ultimately of obstructive pulmonary disease (Stern et al., 2007). Further, the question of whether asthma in survivors of BPD is more likely to be non-atopic is still open, although there is some evidence supporting this hypothesis (Verhaeghe et al., 1990, Ronkainen et al., 2016). In this context, the fact that this signaling pathway is nonetheless implicated as a mediator of gene expression changes in infants with BPD is intriguing and may shed light on the mechanisms involved in the disease. Perhaps it simply reflects airway hyperreactivity in general, but there is little specific evidence supporting this possibility. Further exploration of this pathway’s role in the development of BPD is therefore warranted.

### Genetic interaction data confirms the identification of mediating pathways

Finding specific examples consistent with existing knowledge provides anecdotal evidence that an approach is effective, but more systematic evaluation is needed. One indication that the proposed mediating pathways are, at least in some respect, downstream of the disease genes, would be to identify an excess of epistatic relationships between them. For example, if a mediating pathway looks like that shown in Figure 1, one might expect a higher likelihood of certain kinds of genetic interactions between a disease gene *d* in set *D* and a mediating gene *m* from set *M* than between *d* and genes that are not in a mediating pathway for that disease. The genetic interactions of most interest would be “alleviating” or positive genetic interactions, where the deleterious effect of the double mutant of both *d* and *m* is less severe than would be predicted by combining the independent effects of individual mutations in *d* or *m*. Such relationships might arise when *m* is part of a pathway mediating the response of *d*.

Because few such relationships have yet been characterized in humans or other mammals, we assembled a set of putative positive genetic relationships from orthologous genes in yeast, worm, and fly. For a mediating gene set or pathway *S* and disease gene set *D*, we determine *k*, the number of such relationships between genes in *D* and *S*. We then use a separate permutation test (see Figure 3 and Methods) to assess the probability of finding *k* such relationships with a random gene set of size |*S*|, which we call *p_med_*(*S*).

**Figure 3.**
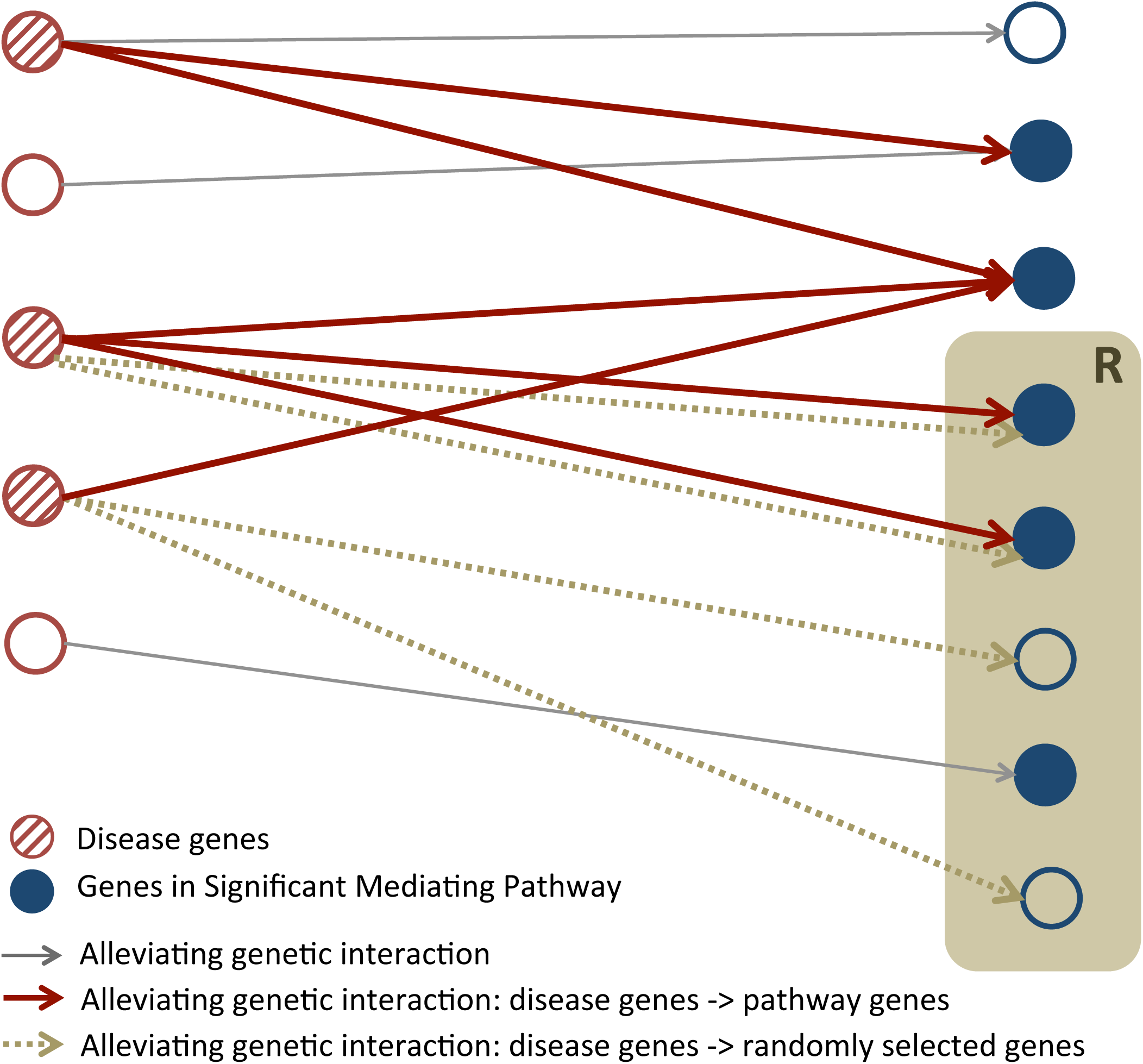
Systematic confirmation of significant mediating pathways. If identified pathways are truly mediating a disease response, the pathways are likely to be downstream of the corresponding disease genes. This relationship can be indicated by an excess of epistatic interactions between disease genes and pathway genes. We count alleviating genetic interactions between disease genes and our identified mediating pathways, and then assess significance by calculating the probability that a gene set of equal size has the same number of alleviating genetic interactions with the disease genes. The null distribution is learned from 10,000 random samples (R) drawn from a pool of downstream genes of any known alleviating genetic interactions.

Overall correlation between *p_cent_*(*S*) and *p_med_*(*S*) varies considerably across the data sets and annotation sets, with a range of 0.07 to 0.74. The lowest correlations come from the Down syndrome data, where this approach is more problematic due to the large number of “disease” (i.e., trisomic) genes that are likely not directly involved in the etiology of the DS phenotype. Instead of overall correlation, however, our goal is to assess the probability that a pathway S having a low *p_cent_*(*S*) value is more likely to have a low *p_med_*(*S*) value as well. We therefore split the range of *p_cent_* values up into deciles, and plotted the probability of finding a *p_med_* score below 0.05 in each decile. A sample plot of these probabilities is shown in Figure 4; the rest are available as supplemental data. If the slope of a linear regression line through the points (the blue line in Figure 4) is significantly less than zero, it suggests that the probability of being a mediator is highest for the most significantly central pathways.

**Figure 4.**
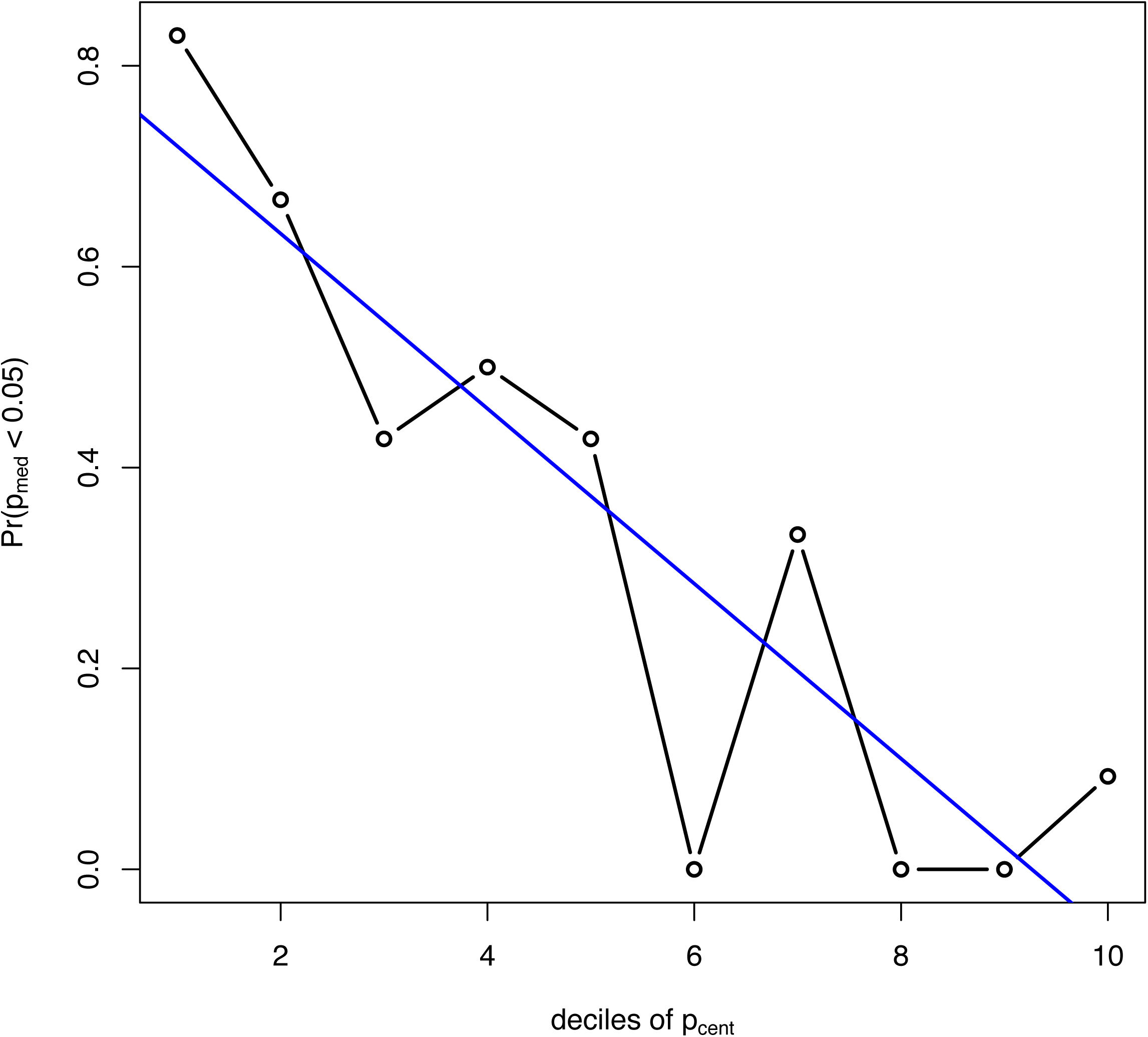
Relationship between p_cent_ and p_med_, for KEGG pathways and COPD. We claim that pathways with low p_cent_ also have low p_med_ values. That is, there is a low probability (p_med_) that a gene set of equal size has as many alleviating genetic interactions with COPD genes as a significant mediating pathway (one with low p_cent_). This plot supports this claim because the linear regression line (in blue) has a slope significantly less than zero (p < 0.0006), suggesting a strong relationship between p_cent_ and p_med_.

Table 3 shows the p-values assessing this property (as reported by the R generalized linear model function glm()). These p-values are consistently high for the DS data, which makes sense for the reason noted above. The test is otherwise significant for the KEGG pathways and for several other pathway/disease combinations. In some of the cases where it is not, such as BPD with the MP gene sets, the first decile does in fact have the highest probability; the poor significance and relatively flat slope occur because there also happen to be many genetic relationships for *p_cent_* values above 0.8.

**Table 3.**
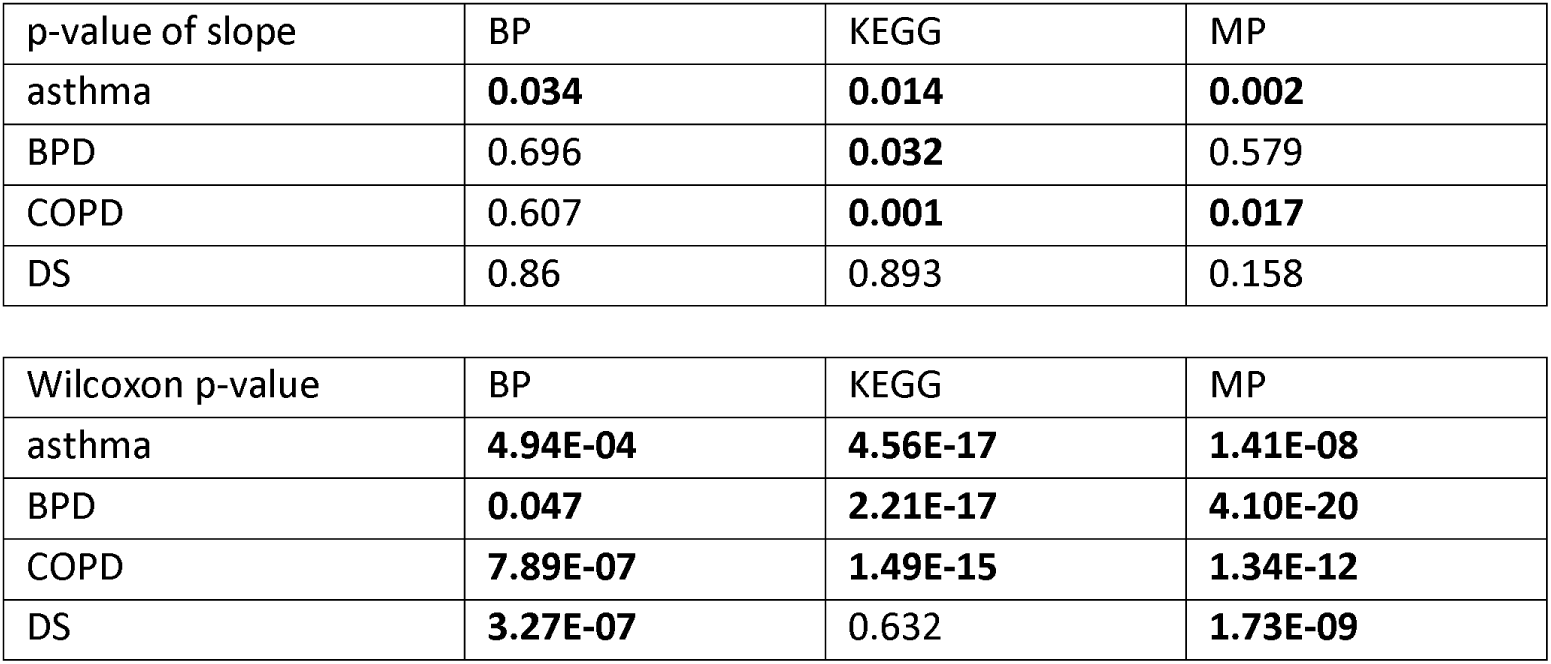
Significance of relationships between *p_cent_* and *p_med_*. The first set of numbers reflects the raw p-values reported by the glm() function in R showing that a regression line fit to the probabilities of having a *p_med_* <0.05 for each decile of *p_cent_* values has a significantly negative slope. The second set shows the raw p-values for Wilcoxon tests comparing the *p_med_* values in just the first decile to those in the remaining deciles combined. Raw p-values below 0.05 appear in bold.

To focus on just the most-significant pathways, we can simply compare the distribution of *p_med_* values in the first bin to that in the remaining bins using a one-sided, non-parametric Wilcoxon test. These significance values are also shown in Table 3. This less strict test is significant in all the pulmonary disease cases and almost all the DS ones as well. Both the tests and the plots support the conclusion that there is enrichment of alleviating genetic relationships between disease genes and pathway genes for the pathways whose *p_cent_* values are in the most significant decile.

Although there are many issues with the data used to produce these results, the conservation of genetic interactions across species being one of the most salient, the trends shown here add systematic evidence to the anecdotal evidence above that this approach can indeed find pathways playing a mediating role in disease processes, potentially leading to the discovery of novel therapeutic interventions. Such an approach may be applied in the context of any disease or phenotype of interest.

## Author Contributions

J.P. and D.K.S. conceived the pathway centrality algorithm and wrote the manuscript. J.P., B.J.H, and D.K.S. designed the statistical method to assess significance of our findings and edited the manuscript. J.P. implemented the algorithm and ran the experiments. J.P. and D.K.S. analyzed the data and provided biological interpretation of the results. All authors read and approved the final version of the manuscript.

## Acknowledgments

We gratefully acknowledge the support of the Eunice Kennedy Shriver National Institute of Child Health & Human Development under award R01HD076140. We thank Diana Bianchi and Jill Maron for helpful discussions. We also thank members of the Tufts Bioinformatics and Computational Biology group for feedback and comments on earlier versions of this work.

## Supplemental Figure Legends

Figure S1.

**Venn diagrams for disease gene sets and differentially expressed genes.** A) Overlap between disease genes for three pulmonary diseases and genes on chromosome 21. B) Overlap between differentially expressed genes in three pulmonary diseases and Down syndrome.

Figure S2.

**R-plots showing that pathways with low p_cent_ also have low p_med_ with linear regression line.** The plots comparable to that in Figure 4 for all combinations of disease and pathway collections are included here.

## Methods and Resources

### Disease-related genes

Disease genes for Asthma and COPD were collected from recent reviews, (Bijanzadeh et al., 2011) and (Bosse, 2012), respectively. A set of 361 Chromosome 21 genes was downloaded from the Molecular Signature DataBase (MSigDB) (Liberzon et al., 2011). We collected genes associated with BPD from Online Mendelian Inheritance in Man (OMIM) (Amberger et al., 2009) and Genopedia (Yu et al., 2010), as described in (Park et al., 2014). All datasets were downloaded on February 3, 2016.

Microarray gene expression profiles for Asthma and BPD were downloaded from the GEO database (accession numbers GSE4302 and GSE32472, respectively). The first measured differential expression in airway epithelial cells between healthy controls and asthma patients (Woodruff et al., 2007), while the second examined expression in peripheral blood cells from infants born preterm with or without BPD (Pietrzyk et al., 2013). From this study, we used only samples taken on postnatal day 5. We selected as differentially expressed genes between disease and control groups those with an adjusted Benjamini-Hochberg t-test p-value below 0.005, yielding 52 and 274 expression-related genes in asthma and BPD, respectively.

For COPD, we downloaded RNA-seq EdgeR results comparing expression in lung cells from COPD patients and controls (GSE57148) (Kim et al., 2015). We identified 266 significantly expressed genes with an EdgeR q-value below 10^-10^. For the control case, we chose a prior study from our group (GSE16176) (Slonim et al., 2009) that compared gene expression in amniotic fluid from second trimester fetuses with Down syndrome to those of age- and sex-matched controls. Using the same criteria as for the other Affymetrix microarray studies (all three of which used the U133 Plus 2.0 arrays) but paired t-tests to deal with the matched samples identified 129 differentially expressed genes in Down syndrome.

Note that there may be disease-associated genes that are also differentially expressed in that disease. To avoid confusion about how to use these in computing pathway centrality, we removed genes from the disease gene sets that also appeared in the corresponding set of differentially expressed genes. The resulting disease gene sets contained 107 (asthma), 73 (BPD), 181 (COPD), and 360 (DS). Supplementary Figure S1 shows the overlaps between these four disease gene sets and the four differentially-expressed gene sets.

### Functional pathways and gene sets

Both the Gene Ontology and the KEGG gene set collections were downloaded from the Molecular Signature DataBase (MSigDB) (Liberzon et al., 2011) on August 18, 2015 (http://software.broadinstitute.org/gsea/msigdb). This GO gene set collection includes 825 biological process terms and 6,178 genes, and the KEGG collection includes 186 pathways and 5,267 genes. In addition, we created a collection of gene sets using data from the Mouse Genome Informatics site (http://www.informatics.jax.org), which includes a database characterizing mutant mouse strains with respect to the Mammalian Phenotype Ontology (Blake et al., 2017). To create gene sets with mammalian phenotype (MP) data, we downloaded the files HMD_HumanPhenotype.rpt and MGI_GenePheno.rpt on February 21, 2016. The first file includes human / mouse orthology mapping and some MP data, while the second includes more MP data characterizing mouse gene mutant strains (other than conditional mutations), but no orthology information. We therefore used the human-mouse orthologs from the first file and then combined the phenotypes from both files, excluding the two normal phenotype labels MP:0002169 (“no abnormal phenotype detected”) and MP:0002873 (“normal phenotype”). We then created for each phenotype a list of genes with at least one mutant allele with that phenotype. These lists were used as functional gene sets, which we describe as the “mammalian phenotype” gene set collection. There are 8,225 phenotypes in the mammalian phenotype data and 7,778 genes.

### Protein-protein and genetic interaction networks

We use two biological networks in our experiments. To measure pathway centrality, physical protein-protein interactions were collected from the Human Integrated Protein-Protein Interaction rEference (HIPPIE) (Alanis-Lobato et al., 2017) database. HIPPIE contains experimentally verified protein interactions with confidence scores. We downloaded the interaction data (version 2) on July 18, 2016 and selected only those interactions described as “high confidence” (≥ 0.73), as these interactions are supported by more reliable evidence. We worked with the largest connected component extracted from the network, which contains 43,475 interactions between 9,379 proteins.

To compute *p_med_* scores based on genetic interaction data, we looked for genetic interaction data featuring alleviating (positive) genetic and phenotypic suppression interactions. Because relatively few of these types of genetic interactions are known for humans, we collected such genetic interactions from *Saccharomyces cerevisiae*, *Drosophila melanogaster*, and *Caenorhabditis elegans.* These interactions came from BioGRID (Stark et al., 2006) (version 3.4.141, downloaded October 14, 2016); the Saccharomyces Genome Database (SGD project, http://www.yeastgenome.org; downloaded October 10, 2016), and Flybase (Attrill et al., 2016) (downloaded August 15, 2016). To find human homologous interaction pairs, we use a mapping downloaded from the HomoloGene database (NCBI Resource Coordinators, 2016) on July 19, 2016. This approach gave us 4,536 pairs of putative positive genetic interactions in humans.

### Identification of disease-mediating pathways

*Pathway centrality* measures the amount of information a set of pathway genes handles by counting paths linking disease genes and differentially expressed genes. While the classical definition of fractional betweenness (FB) for a node *v* is the fraction of shortest paths between all pairs of nodes in a network passing through *v*, our pathway centrality (PC) score will be based on a modified FB score (FB’), which for node *v*, only reflects the shortest paths between disease genes, *D(d)*, and differentially expressed genes, *E(d)*, that passes through *v*. Formally, this is defined as:

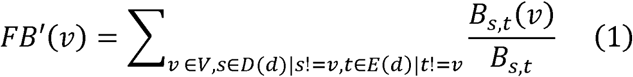

where *V* is the set of all vertices in the protein-protein interaction network, *D(d)* is the set of genes in *V* associated with disease *d*, *E(d)* is the set of differentially expressed genes in *V* for disease *d*, *B*_*s*,*t*_ is the total number of shortest paths between *s* and *t*, and *Bs,t(v)* is the total number of shortest paths between *s* and *t* that pass through *v*.

We define *pathway centrality* as the average fractional betweenness score across all genes in a pathway, counting only the shortest paths from disease genes to differentially expressed genes. Specifically, for a pathway *k* containing the gene set *P(k)*, pathway centrality(k) is defined as:

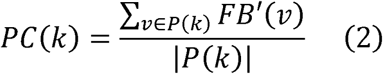

### Assessing significance of identified mediating pathways: p_*cent*_

The significance of the observed *pathway centrality* score of a pathway or gene set S is assessed using a null distribution learned from a permutation with 10,000 random gene sets of size |S|. We further require that the random gene sets be drawn from a collection of genes satisfying two conditions: they should belong to at least one pathway, and should have degree ≥ 2. The reason for these restrictions is to reduce biases that artificially reduce the *p_cent_* values, making the majority of pathways appear to be significantly central. Pathway genes are known to be relatively central in protein interaction networks, and therefore, it is likely that pathway genes have higher fractional betweenness than those that are not involved in well-annotated functional processes.

Further, to the extent that protein interaction networks are scale-free, random samples will likely be enriched for genes of degree 1 if we do not somehow force the random sampling process to select higher degree genes. But this is problematic because the distribution of *pathway centrality* from permutation tests may then be left-skewed, given that the various centrality measurements tend to be correlated (Goh et al., 2003). Thus, to systematically exclude low-degree genes from permutation, we decided to use only genes in the 2-core of the protein-protein interaction network. A *k*-*core* of a network is a maximal group of nodes, all of which are connected to at least *k* other nodes in the network (Bhawalkar et al., 2012). This approach improves the observed distribution of *p_cent_* values. It may also reduce the significance of pathways containing mostly low-degree genes, but this applies to relatively few of the gene sets in the considered collections.

### Evaluation of identified disease-mediating pathways

We compute *p_med_* by counting the number of phenotypic alleviation or suppression interactions known to exist between disease genes and the mediators. To assess how surprising it is to see the observed number of such interactions, we compute a null distribution of such counts between the *same* set of disease genes and 10,000 random gene sets of equal size as the mediating pathway. Again we impose restrictions on the source of random genes to avoid over-estimating significance. Here, random gene sets are drawn from a pool of genes that belong to at least one pathway in the collection and that are downstream genes of any alleviating genetic or phenotypic suppression interactions.

## Supplemental Information

We are providing one excel file containing our full experimental results, two pdf files containing two Venn diagrams and nine R plots, and three cytoscape session files.

## REFERENCES

Abman, S. H., Schaffer, M. S., Wiggins, J., Washington, R., Manco-Johnson, M. & Wolfe, R.R. 1987. Pulmonary vascular extraction of circulating norepinephrine in infants with bronchopulmonary dysplasia. Pediatr Pulmonol, 3, 386–91.

Alanis-Lobato, G., Andrade-Navarro, M. A. & Schaefer, M. H. 2017. Hippie v2.0: enhancing meaningfulness and reliability of protein-protein interaction networks. Nucleic Acids Res, 45, D408–D414.

Ali, S., Hirschfeld, A. F., Mayer, M. L., Fortuno, E. S., 3RD, Corbett, N., Kaplan, M., Wang, S., Schneiderman, J., Fjell, C. D., Yan, J., Akhabir, L., Aminuddin, F., Marr, N., Lacaze-Masmonteil, T., Hegele, R. G., Becker, A., Chan-Yeung, M., Hancock, R. E., Kollmann, T. R., Daley, D., Sandford, A. J., Lavoie, P. M. & Turvey, S. E. 2013. Functional genetic variation in NFKBIA and susceptibility to childhood asthma, bronchiolitis, and bronchopulmonary dysplasia. J Immunol, 190, 3949–58.

Amberger, J., Bocchini, C. A., Scott, A. F. & Hamosh, A. 2009. McKusick’s Online Mendelian Inheritance in Man (OMIM). Nucleic Acids Res, 37, D793–6.

Apostolou, A., Kerenidi, T., Michopoulos, A., Gourgoulianis, K. I., Noutsias, M., Germenis, A. E. & Speletas, M. 2016. Association between TLR2/TLR4 gene polymorphisms and COPD phenotype in a Greek cohort. Herz.

Ashburner, M., Ball, C. A., Blake, J. A., Botstein, D., Butler, H., Cherry, J. M., Davis, A. P., Dolinski, K., Dwight, S. S., Eppig, J. T., Harris, M. A., Hill, D. P., Issel-Tarver, L., Kasarskis, A., Lewis, S., Matese, J. C., Richardson, J. E., Ringwald, M., Rubin, G. M. & Sherlock, G. 2000. Gene ontology: tool for the unification of biology. The Gene Ontology Consortium. Nat Genet, 25, 25–9.

Attrill, H., Falls, K., Goodman, J. L., Millburn, G. H., Antonazzo, G., Rey, A. J., Marygold, S. J. & Flybase, C. 2016. FlyBase: establishing a Gene Group resource for Drosophila melanogaster. Nucleic Acids Res, 44, D786–92.

Barnes, P. J. 2016. Kinases as Novel Therapeutic Targets in Asthma and Chronic Obstructive Pulmonary Disease. Pharmacol Rev, 68, 788–815.

Barnes, P. J., Drazen, J. M., Rennard, S. I. & Thomson, N. C. 2009. Asthma and COPD Basic Mechanisms and Clinical Management Second Edition Preface to the 2nd Edition. Asthma and Copd: Basic Mechanisms and Clinical Management, 2nd Edition, pp. 178ff.

Belcastro, R., Lopez, L., Li, J., Masood, A. & Tanswell, A. K. 2015. Chronic lung injury in the neonatal rat: up-regulation of TGFbeta1 and nitration of IGF-R1 by peroxynitrite as likely contributors to impaired alveologenesis. Free Radic Biol Med, 80, 1–11.

Bhawalkar, K., Kleinberg, J., Lewi, K., Roughgarden, T. & Sharma, A. 2012. Preventing Unraveling in Social Networks: The Anchored k-Core Problem. Automata, Languages, and Programming, Icalp 2012, Pt Ii, 7392, 440–451.

Biggadike, K., Ahmed, M., Ball, D. I., Coe, D. M., Dalmas Wilk, D. A., Edwards, C. D., Gibbon, B. H., Hardy, C. J., Hermitage, S. A., Hessey, J. O., Hillegas, A. E., Hughes, S. C., Lazarides, L., Lewell, X. Q., Lucas, A., Mallett, D. N., Price, M. A., Priest, F. M., Quint, D. J., Shah, P., Sitaram, A., Smith, S. A., Stocker, R., Trivedi, N. A., Tsitoura, D. C. & Weller, V. 2016. Discovery of 6-Amino-2-{[(1S)-1-methylbutyl]oxy}-9-[5-(1-piperidinyl)pentyl]-7,9-dihydro-8H-pu rin-8-one (GSK2245035), a Highly Potent and Selective Intranasal Toll-Like Receptor 7 Agonist for the Treatment of Asthma. J Med Chem, 59, 1711–26.

Bijanzadeh, M., Mahesh, P. A. & Ramachandra, N. B. 2011. An understanding of the genetic basis of asthma. Indian J Med Res, 134, 149–61.

Blake, J. A., Eppig, J. T., Kadin, J. A., Richardson, J. E., Smith, C. L., Bult, C. J. & The Mouse Genome Database, G. 2017. Mouse Genome Database (MGD)-2017: community knowledge resource for the laboratory mouse. Nucleic Acids Res, 45, D723–D729.

Bodas, M., Tran, I. & Vij, N. 2012. Therapeutic strategies to correct proteostasis-imbalance in chronic obstructive lung diseases. Curr Mol Med, 12, 807–14.

Bosse, Y. 2012. Updates on the COPD gene list. Int J Chron Obstruct Pulmon Dis, 7, 607–31.

Bouchecareilh, M. & Balch, W. E. 2012. Proteostasis, an emerging therapeutic paradigm for managing inflammatory airway stress disease. Curr Mol Med, 12, 815–26.

Buckley, S., Barsky, L., Weinberg, K. & Warburton, D. 2005. In vivo inosine protects alveolar epithelial type 2 cells against hyperoxia-induced DNA damage through MAP kinase signaling. Am J Physiol Lung Cell Mol Physiol, 288, L569–75.

Bult, C. J., Eppig, J. T., Blake, J. A., Kadin, J. A., Richardson, J. E. & Mouse Genome Database, G. 2016. Mouse genome database 2016. Nucleic Acids Res, 44, D840–7.

Carrera, P., Di Resta, C., Volonteri, C., Castiglioni, E., Bonfiglio, S., Lazarevic, D., Cittaro, D., Stupka, E., Ferrari, M., Somaschini, M., BPD & Genetics Study, G. 2015. Exome sequencing and pathway analysis for identification of genetic variability relevant for bronchopulmonary dysplasia (BPD) in preterm newborns: A pilot study. Clin Chim Acta, 451, 39–45.

Chen, Z. H., Kim, H. P., Ryter, S. W. & Choi, A. M. 2008. Identifying targets for COPD treatment through gene expression analyses. Int J Chron Obstruct Pulmon Dis, 3, 359–70.

Chetty, A., Cao, G. J. & Nielsen, H. C. 2006. Insulin-like Growth Factor-I signaling mechanisms, type I collagen and alpha smooth muscle actin in human fetal lung fibroblasts. Pediatr Res, 60, 389–94.

Delude, C. M. 2015. Deep phenotyping: The details of disease. Nature, 527, S14–5.

Erdos, D. 2015. Centrality measures and analyzing dot-product graphs. Boston: Boston University.

Everett, M. G. & Borgatti, S. P. 1999. The centrality of groups and classes. Journal of Mathematical Sociology, 23, 181–201.

Faura Tellez, G., Willemse, B. W., Brouwer, U., Nijboer-Brinksma, S., Vandepoele, K., Noordhoek, J. A., Heijink, I., DE Vries, M., Smithers, N. P., Postma, D. S., Timens, W., Wiffen, L., Van Roy, F., Holloway, J. W., Lackie, P. M., Nawijn, M. C. & Koppelman, G. H. 2016. Protocadherin-1 Localization and Cell-Adhesion Function in Airway Epithelial Cells in Asthma. PLoS One, 11, e0163967.

Fox, A. D., Hescott, B. J., Blumer, A. C. & Slonim, D. K. 2011. Connectedness of PPI network neighborhoods identifies regulatory hub proteins. Bioinformatics, 27, 1135–42.

Freeman, L. C., Borgatti, S. P. & White, D. R. 1991. Centrality in Valued Graphs - a Measure of Betweenness Based on Network Flow. Social Networks, 13, 141–154.

Gagliardo, R., Chanez, P., Profita, M., Bonanno, A., Albano, G. D., Montalbano, A. M., Pompeo, F., Gagliardo, C., Merendino, A. M. & Gjomarkaj, M. 2011. IkappaB kinase-driven nuclear factor-kappaB activation in patients with asthma and chronic obstructive pulmonary disease. J Allergy Clin Immunol, 128, 635–45 e1-2.

Goh, K. I., Oh, E., Kahng, B. & Kim, D. 2003. Betweenness centrality correlation in social networks. Phys Rev E Stat Nonlin Soft Matter Phys, 67, 017101.

Gould, H. J. & Sutton, B. J. 2008. IgE in allergy and asthma today. Nat Rev Immunol, 8, 205–17.

Hudson, N. J., Dalrymple, B. P. & Reverter, A. 2012. Beyond differential expression: the quest for causal mutations and effector molecules. BMC Genomics, 13, 356.

Ishii, Y. 1999. [Role of adhesion molecules in the pathogenesis of COPD]. Nihon Rinsho, 57, 1965–71.

Kanehisa, M., Sato, Y., Kawashima, M., Furumichi, M. & Tanabe, M. 2016. KEGG as a reference resource for gene and protein annotation. Nucleic Acids Res, 44, D457–62.

Kim, E. K. & Choi, E. J. 2010. Pathological roles of MAPK signaling pathways in human diseases. Biochim Biophys Acta, 1802, 396–405.

Kim, W. J., Lim, J. H., Lee, J. S., Lee, S. D., Kim, J. H. & Oh, Y. M. 2015. Comprehensive Analysis of Transcriptome Sequencing Data in the Lung Tissues of COPD Subjects. Int J Genomics, 2015, 206937.

Kim, Y.-A., Wuchty, S. & Przytycka, T. M. 2011. Identifying Causal Genes and Dysregulated Pathways in Complex Diseases. PLOS Computational Biology, 7, el001095.

Kwinta, P., Lis, G., Klimek, M., Grudzien, A., Tomasik, T., Poplawska, K. & Pietrzyk, J. J. 2013. The prevalence and risk factors of allergic and respiratory symptoms in a regional cohort of extremely low birth weight children (<1000 g). Ital J Pediatr, 39, 4.

Liberzon, A., Subramanian, A., Pinchback, R., Thorvaldsdottir, H., Tamayo, P. & Mesirov, J. P. 2011. Molecular signatures database (MSigDB) 3.0. Bioinformatics, 27, 1739–40.

Lofqvist, C., Hellgren, G., Niklasson, A., Engstrom, E., Ley, D., Hansen-Pupp, I. & Consortium, W. 2012. Low postnatal serum IGF-I levels are associated with bronchopulmonary dysplasia (BPD). Acta Paediatr, 101, 1211–6.

Malash, A. H., Ali, A. A., Samy, R. M. & Shamma, R. A. 2016. Association of TLR polymorphisms with bronchopulmonary dysplasia. Gene, 592, 23–8.

Mason, O. & Verwoerd, M. 2007. Graph theory and networks in Biology. IET Syst Biol, 1, 89–119.

Missiuro, P. V., Liu, K., Zou, L., Ross, B. C., Zhao, G., Liu, J. S. & Ge, H. 2009. Information flow analysis of interactome networks. PLoS Comput Biol, 5, el000350.

Nabieva, E., Jim, K., Agarwal, A., Chazelle, B. & Singh, M. 2005. Whole-proteome prediction of protein function via graph-theoretic analysis of interaction maps. Bioinformatics, 21 Suppl 1, i302–10.

Ncbi Resource Coordinators 2016. Database resources of the National Center for Biotechnology Information. Nucleic Acids Res, 44, D7–19.

O’shea, J. J., Schwartz, D. M., Villarino, A. V., Gadina, M., Mcinnes, I. B. & Laurence, A. 2015. The JAK-STAT pathway: impact on human disease and therapeutic intervention. Annu Rev Med, 66, 311–28.

Park, J., Wick, H. C., Kee, D. E., Noto, K., Maron, J. L. & Slonim, D. K. 2014. Finding novel molecular connections between developmental processes and disease. PLoS Comput Biol, 10, el003578.

Petrikin, J. E., Gaedigk, R., Leeder, J. S. & Truog, W. E. 2010. Selective Toll--like receptor expression in human fetal lung. Pediatr Res, 68, 335–8.

Pietrzyk, J. J., Kwinta, P., Wollen, E. J., Bik-Multanowski, M., Madetko-Talowska, A., Gunther, C. C., Jagla, M., Tomasik, T. & Saugstad, O. D. 2013. Gene expression profiling in preterm infants: new aspects of bronchopulmonary dysplasia development. PLoS One, 8, e78585.

Ramsay, P. L., O’brian Smith, E., Hegemier, S. & Welty, S. E. 1998. Early clinical markers for the development of bronchopulmonary dysplasia: soluble E-Selectin and ICAM-1. Pediatrics, 102, 927–32.

Ronkainen, E., Kaukola, T., Marttila, R., Hallman, M. & Dunder, T. 2016. School-age children enjoyed good respiratory health and fewer allergies despite having lung disease after preterm birth. Acta Paediatr, 105, 1298–1304.

Sackmann, E. K., Berthier, E., Schwantes, E. A., Fichtinger, P. S., Evans, M. D., Dziadzio, L. L., Huttenlocher, A., Mathur, S. K. & Beebe, D. J. 2014. Characterizing asthma from a drop of blood using neutrophil chemotaxis. Proc Natl Acad Sci U S A, 111, 5813–8.

Sakurai, R., Villarreal, P., Husain, S., Liu, J., Sakurai, T., Tou, E., Torday, J. S. & Rehan, V. K. 2013. Curcumin protects the developing lung against longterm hyperoxic injury. Am J Physiol Lung Cell Mol Physiol, 305, L301–11.

Sharma, V., Michel, S., Gaertner, V., Franke, A., Vogelberg, C., Von Berg, A., Bufe, A., Heinzmann, A., Laub, O., Rietschel, E., Simma, B., Frischer, T., Genuneit, J., Potaczek, D. P. & Kabesch, M. 2014. A role of FCER1A and FCER2 polymorphisms in IgE regulation. Allergy, 69, 231–6.

Shaykhiev, R. & Crystal, R. G. 2013. Innate immunity and chronic obstructive pulmonary disease: a mini-review. Gerontology, 59, 481–9.

Siltanen, M., Kajosaari, M., Pohjavuori, M. & Savilahti, E. 2001. Prematurity at birth reduces the long-term risk of atopy. J Allergy Clin Immunol, 107, 229–34.

Simon, A. R., Takahashi, S., Severgnini, M., Fanburg, B. L. & Cochran, B. H. 2002. Role of the JAK-STAT pathway in PDGF-stimulated proliferation of human airway smooth muscle cells. Am J Physiol Lung Cell Mol Physiol, 282, L1296–304.

Slonim, D. K., Koide, K., Johnson, K. L., Tantravahi, U., Cowan, J. M., Jarrah, Z. & Bianchi, D. W. 2009. Functional genomic analysis of amniotic fluid cell-free mRNA suggests that oxidative stress is significant in Down syndrome fetuses. Proc Natl Acad Sci U S A, 106, 9425–9.

Stark, C., Breitkreutz, B. J., Reguly, T., Boucher, L., Breitkreutz, A. & Tyers, M. 2006. BioGRID: a general repository for interaction datasets. Nucleic Acids Res, 34, D535–9.

Stern, D. A., Morgan, W. J., Wright, A. L., Guerra, S. & Martinez, F. D. 2007. Poor airway function in early infancy and lung function by age 22 years: a non-selective longitudinal cohort study. Lancet, 370, 758–64.

Suthram, S., Beyer, A., Karp, R. M., Eldar, Y. & Ideker, T. 2008. eQED: an efficient method for interpreting eQTL associations using protein networks. Molecular Systems Biology, 4.

Suzuki, M., Makita, H., Konno, S., Shimizu, K., Kimura, H., Kimura, H., Nishimura, M. & Hokkaido, C. C. S. I. 2016. Asthma-like Features and Clinical Course of Chronic Obstructive Pulmonary Disease. An Analysis from the Hokkaido COPD Cohort Study. Am J Respir Crit Care Med, 194, 1358–1365.

Tu, Z., Wang, L., Arbeitman, M. N., Chen, T. & Sun, F. 2006. An integrative approach for causal gene identification and gene regulatory pathway inference. Bioinformatics, 22, e489–e496.

Vale, K. 2016. Targeting the JAK-STAT pathway in the treatment of ‘Th2-high’ severe asthma. Future Med Chem, 8, 405–19.

Verhaeghe, M., De Wolf, M., Lagrou, A., Van Dessel, G., Hilderson, H. & Dierick, W. 1990. Identification of essential amino acids in the active center of thyroidal NAD+ glycohydrolase. Int J Biochem, 22, 197–202.

Weltman, J. K. & Karim, A. S. 1998. Interleukin-5: a proeosinophil cytokine mediator of inflammation in asthma and a target for antisense therapy. Allergy Asthma Proc, 19, 257–61.

Wong, P. M., Lees, A. N., Louw, J., Lee, F. Y., French, N., Gain, K., Murray, C. P., Wilson, A. & Chambers, D. C. 2008. Emphysema in young adult survivors of moderate-to-severe bronchopulmonary dysplasia. Eur Respir J, 32, 321–8.

Woodruff, P. G., Boushey, H. A., Dolganov, G. M., Barker, C. S., Yang, Y. H., Donnelly, S., Ellwanger, A., Sidhu, S. S., Dao-Pick, T. P., Pantoja, C., Erle, D. J., Yamamoto, K. R. & Fahy, J. V. 2007. Genome-wide profiling identifies epithelial cell genes associated with asthma and with treatment response to corticosteroids. Proc Natl Acad Sci U S A, 104, 15858–63.

Woodside, D. G. & Vanderslice, P. 2008. Cell adhesion antagonists: therapeutic potential in asthma and chronic obstructive pulmonary disease. BioDrugs, 22, 85–100.

Xu, D., Guthrie, J. R., Mabry, S., Sack, T. M. & Truog, W. E. 2006. Mitochondrial aldehyde dehydrogenase attenuates hyperoxia-induced cell death through activation of ERK/MAPK and PI3K-Akt pathways in lung epithelial cells. Am J Physiol Lung Cell Mol Physiol, 291, L966–75.

Yeger-Lotem, E., Riva, L., Su, L. J., Gitler, A. D., Cashikar, A. G., King, O. D., Auluck, P. K., Geddie, M. L., Valastyan, J. S. & Karger, D. R. 2009. Bridging high-throughput genetic and transcriptional data reveals cellular responses to alpha-synuclein toxicity. Nature genetics, 41, 316–323.

Yu, H., Kim, P. M., Sprecher, E., Trifonov, V. & Gerstein, M. 2007. The importance of bottlenecks in protein networks: correlation with gene essentiality and expression dynamics. PLoS Comput Biol, 3, e59.

Yu, H., Lin, M., Wang, X., Wang, S. & Wang, Z. 2016. Toll-like receptor 4 polymorphism is associated with increased susceptibility to chronic obstructive pulmonary disease in Han Chinese patients with chronic periodontitis. J Oral Sci, 58, 555–560.

Yu, W., Clyne, M., Khoury, M. J. & Gwinn, M. 2010. Phenopedia and Genopedia: disease-centered and gene-centered views of the evolving knowledge of human genetic associations. Bioinformatics, 26, 145–6.

Zhou, L. & Hershenson, M. B. 2003. Mitogenic signaling pathways in airway smooth muscle. Respir Physiol Neurobiol, 137, 295–308.

